# SpeciefAI: Multi-species mRNA-level Antibody Framework Generation using Transformers

**DOI:** 10.64898/2026.03.16.712018

**Authors:** Dominik Grabarczyk, Mikołaj Kocikowski, Maciej Parys, Shay B. Cohen, Javier Antonio Alfaro

## Abstract

**Motivation:** Encoding antibodies (Abs) and nanobodies (Nbs) as mRNA enables in vivo production of therapeutic proteins. However, this approach requires meeting two species-dependent requirements: the mRNA encoding must support efficient expression in the host species, and the encoded protein sequence must resemble the natural Ab repertoire of the recipient species to minimize immunogenicity. These requirements motivate species-conditioned generative models for joint mRNA and protein design.

**Results:** We propose SpeciefAI a transformer-based model for multi-species Ab and Nb species sequence-harmonisation by generation of novel Framework Regions (FRs) tailored to input Complementarity-Determining Regions (CDRs). Our model works directly in the mRNA space and learns the correspondence between FRs and CDRs in six species. The model is capable of generating sequences with a highly similar distribution to natural sequences and a mean absolute difference in codon adaptation index (CAI) of 0.013 and 0.033 for humans and dogs respectively. We show that the generated human sequences are highly human (0.95 T20 score) and canine sequences highly canine (0.95 cT20 score). We furthermore demonstrate that we can generate diverse candidate sequences using our method.

**Availability and Implementation:** Source code is available on https://github.com/Dominko/SpeciefAI. OAS and COGNANO data are publicly available on https://opig.stats.ox.ac.uk/webapps/oas/ and https://cognanous.com/datasets/vhh-corpus (preprocessed versions available upon request). Canine data is available on https://zenodo.org/records/18301526.

## Introduction

Antibodies (Abs) are specialized antigen-binding proteins of the adaptive immune system. Individually, they exhibit high specificity and affinity for their respective targets. As a repertoire, they exhibit remarkable sequence diversity due to their origin in variable–diversity–joining (VDJ) recombination, which enables them to recognize a broad spectrum of antigens [18]. These characteristics make them an attractive molecule format for therapeutic development against cancer, anto-immune, viral, and other diseases.

A typical mammalian Ab is composed of two identical heavy chains and two identical light chains. Each chain can further be subdivided into two sections: a variable domain, called *V*_*H*_ for the heavy chain and *V*_*L*_ for the light chain, and a constant domain.

The combination of one *V*_*H*_ and one *V*_*L*_ forms a single antigen-binding site. Each variable domain consists of three complementarity-determining regions (CDRs), which can bind onto the specific epitope on a target antigen, and four structural frameworks (FRs), which form a scaffold enabling the CDRs to reach their functional conformation [3]. Camelid Abs, originating from species such as camels and alpacas, are unique in that they lack light chains. This architecture means that a single camelid Ab chain can be developed into a binder protein. Reduction from 6 CDRs on 2 chains to 3 CDRs on a single chain drastically simplifies *de novo* protein design and removes the need to model the 3d interactions between chains. These features, along with high stability make camelid VHHs an attractive template for engineering therapeutic single domain antibodies (sdAb) also known as nanobodies (Nb) [2].

One key challenge in using Abs and Nbs as therapeutics is reducing the risk of immune reaction to the foreign proteins being introduced into the patient’s body. While the amino acid (AA) sequences of CDRs primarily depend on the target antigen, FRs sequences are more species-specific. This requires Abs developed based on a model species to first be adjusted to resemble its intended host species before administration [15] to reduce the risk of immunogenicity. While Nbs appear less immunogenic, they are not risk-free [2]. The ability to make such adjustments has utility beyond the development of human therapeutics: some pathogens and antigens are equally relevant in veterinary medicine.

Adapting Abs to new hosts traditionally has been done using methods such as Framework shuffling followed by backtranslation [4]. In framework shuffling, high-throughput in-vitro screening is used to test candidate sequences. Candidate sequences are selected by supplementing CDRs of interest with randomly selected FRs from the target species. As high-throughput is very expensive, recently AI-based computational methods have been developed to address sequence-species harmonization. For example BioPhi [16] uses attention based mutation prediction and HuAbDiffusion uses a transformer based diffusion model for FR generation [11]. These computational methods, however, still operate on protein level, and rely on subsequent backtranslation. These methods also often focus on only humanisation.

Modifications aimed at making the sequences resemble its intended host are also important on the nucleic acid (NA) level. While some therapeutics are manufactured in e.g. *E. coli* bacteria [9], an emerging trend is to produce them *in vivo* in the host as mRNA therapeutics. This method of delivery has potential for significantly decreased costs of Ab therapeutics [14]. The hosts for expression have very different codon adaptation indices (CAIs) and relative tRNA abundance levels, meaning a sequence optimal for one host will not express as effectively in another [20]. To enable effective protein production in new host systems, the pre-optimised AA sequence is commonly back-translated into a NA sequence (mRNA). However, the back-translation method has a significant drawback, as the choice of protein sequence places significant constraints on the NA sequence.

In effect, the FRs of an Ab/Nb mRNA therapeutic must satisfy two species-dependent requirements: the mRNA encoding must support efficient, well-tolerated expression in the host species, and the encoded protein sequence must resemble the natural Ab repertoire of the recipient species to minimize immunogenicity. In humans this is often called humanisation, but to be invariant with respect to the species we perform this on, we will apply the general term species sequence-harmonisation.

To address these requirements for species sequence-harmonisation, we developed SpeciefAI, a T5 transformer-based large language model (LLM) for multi-species Ab species sequence-harmonisation and mRNA optimisation. Our model harmonises the AA sequence by generating species-appropriate FRs based on the input CDRs. Simultaneously, it addresses the problem of mRNA optimisation by generating directly in the mRNA space, rather than generating proteins first. Here, we demonstrate the ability of the model to generate Abs for two clinically relevant species, humans and dogs, and to computationally harmonise sequences with respect to these species. We also show its ability to generate diverse *de novo* sequences for a single set of target CDRs and that the generated sequences are highly similar to the original set of FRs associated with the target CDRs. We also investigate the way the model represents its underlying understanding of Abs.

## Methods

The model used in this project is based on a T5 text-to-text transformer architecture [17]. The model uses a standard encoder-decoder model, since there are clear pairs of inputs and outputs, i.e. CDRs and FRs, where the inputs should provide context for generation throughout the whole output. The model has 12 layers each consisting of 12 heads with a *d*_*model*_ (latent representation dimension) of 1,536. The feed-forward layers each have a dimensionality of 4,096. We used positional encoding to incorporate long range relationships between tokens.

We training our model on mRNA sequence data from several species, an overview of which can be found in Table 1. Human, murine, rat and monkey data comes from the Observed Antibody Space (OAS) dataset [13]. The canine data comes from the work of *Lisowska et al*. [10], which has previously been published as AA sequences [8]. Alpaca data was obtained from the COGNANO VHHCorpus-2M dataset [22]. Alpaca data involves an additional preprocesing step, where the AA data is first back-translated into *E. Coli* mRNA using a high-expression frequency table.

**Table 1.** Overview of number of Ab sequences used in the project per species, data split and origin. *The murine sequences are split between normal murine sequences, and sequences coming from 5 clinically relevant subtypes of mice: HIS mice, BALB/c mice, C57BL/6 mice, RAG2-GFP/129Sve mice and Swiss Webster mice. No distinction is made within the project between these strains.

| Species | Origin | Training Sequences | Validation Sequences | Test Sequences |
| --- | --- | --- | --- | --- |
| Human | OAS | 503,716,618 | 10,000 | 10,000 |
| Mouse* | OAS | 109,019,266 | 10,000 | 10,000 |
| Rat | OAS | 5,052,758 |  |  |
| Monkey | OAS | 3,192,348 |  |  |
| Dog | DoggifAI | 445,672 | 10,000 | 10,000 |
| Alpaca | COGNANO | 1,585,593 | 10,000 | 10,000 |

A balanced held out dataset of 10,000 randomly selected sequences each from the human, murine, canine and alpaca hosts are held out for validation purposes (40,000 in total), and another 10,000 of each as held-out test set.

Each data-point is further preprocessed, by splitting the sequences into CDRs and FRs according to the IMGT numbering scheme using the Abnum web-tool [1]. Sequences which cannot be split using this tool or which are out of frame were discarded. Secondly, each data-point is additionally labelled with its respective species and chain type.

Sequences are tokenised using a 6-mer based tokeniser, such that a token corresponds to a sequence of 6 nucleotides. This approach balances the requirements in terms of vocabulary size (4096 tokens, excluding meta tokens) while preventing any potential frame shifts, by being a multiple of codon triplets.

### Training

The model is trained in two stages, pretraining followed by finetuning. Pretraining is done using semi-supervised learning. Ab sequences have spans of random length and location masked and replaced by placeholder tokens such that 20% of the sequence is masked and the mean length of the spans is 10 tokens. The model is then finetuned by giving the masked Ab sequence as input and the masked spans as desired output.

Each model trained is pretrained on the entire training set using a effective batch size of 1,024 spread out over several gradient accumulation steps. We set a computational limit of at most 24 hours on two NVIDIA GH200 Superchips per model. on the first 10% of the steps. Intermediate perplexity evaluation on the validation set is performed every 1,000 steps. After pretraining on the full training set, the best checkpoint is selected based on the lowest perplexity on the validation set.

The model takes mRNA CDR sequences of an Ab as input, and the FR sequence of the same Ab as desired output. Both are tagged with species and chain identifiers. Since our dataset is highly imbalanced, we rebalance our dataset for finetuning such that under-represented species are oversampled and over-represented species are undersampled. We sample such that the proportion *π*_*y*_ of any class *y* in our set of labels *𝒴* = *{human, dog, mouse, rat, monkey, alpaca}*remains constant in every batch. We rebalance using the following formula, where *n*_*y*_ is the number of instances of class *y* in the dataset and *λ* is a hyper-parameter determining the strength of rebalancing: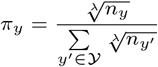. We set *λ* to 4 based on experimental results.

The model is finetuned on the training dataset and the best model/checkpoint are selected based on the validation set.

We define two metrics to score our models. Firstly *E*_*d*_(*s*^*n*^, *s*^*o*^) which is defined as the minimum number of substitutions and gaps between the generated sequence *s*^*n*^ to the original sequence *s*^*o*^ in the set of all possible alignments *A*(*s*^*n*^, *s*^*o*^). This is equivalent to a global pairwise sequence alignment where all mutations and gaps are treated equally. This can be formally expressed as:

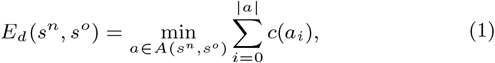

where the cost function *c*(*a*_*i*_) equals 1 if there is a mismatch or gap, and 0 if the positions match.

On protein level, different amino acid substitutions for a given amino acid are not equivalent, due to different chemical properties of amino acids. Additionally, in both mRNA and protein sequences fewer but longer gaps are preferred to many short ones. To address this, we introduce a second, biology-aware error metric *E*_*B*_(*s*^*n*^, *s*^*o*^). In *E*_*B*_(*s*^*n*^, *s*^*o*^), substitutions are scored according to a PAM30 substitution metrics when applied to proteins [5]. Furthermore, gap insertions are penalised by -11 while extensions only by -1. This results in the following:

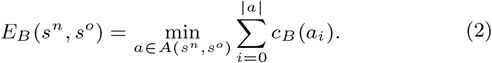

The biology-aware cost function *c*_*B*_(*a*_*i*_) equals 11 if this alignment opens a new gap, 1 if it extends a gap, the PAM30 value of the substitutions if the amino acids are mismatched, and 0 if the amino acids match. The gap insertion and extension penalties are common default parameters, used e.g. by NCBI.

Sequences are decoded from model outputs in two standard ways, greedy decoding and top-*k* sampling. In greedy decoding, the next token *x*_*i*+1_ is the word *v* from the full vocabulary *V* with the maximum likelihood, given the preceding tokens *x*_0_…*x*_*i*_ and the hidden representation *h* of the input sequence. In top-*k* sampling, the next token *x*_*i*+1_ is drawn randomly from the set of tokens *V* ^*k*^ of size *k* with largest probability. Unless otherwise specified, greedy decoding was used in the experiments.

### Multi-species generation properties

To evaluate our model, we sought to investigate the species-specific properties of sequences generated by our multi-species model. Specifically the models ability to match a species’ CAI distribution, and the underlying sequence distribution seen in the test data.

We investigated the ability of our model to match the CAI distribution of the different species. We elected to calculate the CAI per species over the test set sequences, since the distribution may differ from the species’ CAI computed over the whole genome. We compute the difference between the CAI of the *de novo* sequences of each of the test set species and the CAI of the original test set sequences for that species.

We used a T-distributed Stochastic Neighbour Embedding (t-SNE) [12] to obtain a visual representation of the distribution of the original and *de novo* sequences. The 2-dimensional t-SNE is computed over one-hot embedding vectors of each of the original and *de novo* sequences. We then computed a Gaussian Kernel Density Estimation (KDE) of the points from the original and *de novo* datasets using Scott’s Rule to select the bandwidth [19] to allow us to compare their distributions.

Since we finetuned our model as a multi-species model, i.e. on data from multiple species at once, we also sought to compare its performance to that of single species models.

Specifically we compare a model finetuned on the whole dataset using rebalancing to two models finetuned on the clinically relevant human and canine datasets respectively. The latter is especially relevant, since it is the smallest dataset which might get either overfitted or, conversely, be under-represented in the multi-lingual model.

### High-throughput generation

In many practical applications it is beneficial to test many different candidate Ab sequences for a given target. Indeed, the ability to generate a large number of diverse candidate sequences is a core strength of the generative AI approach for Ab species sequence-harmonisation.

It is therefore important to understand the statistical properties of large-scale stochastic sampling from the model for a single CDR. Specifically, we investigate how many unique sequences our model generates and whether these sequences are reasonably close to the original germline sequence.

### Interlingual Ab representation

We sought to investigate whether the model learns an interlingual representation Abs or if it learns to generate FRs independently for each species. To this end, we generated FRs for a single set of CDRs for each of the species in the project.

If the model generates FRs with a clear consensus, it can tell us something about the underlying Ab interlingual representation of an Ab learned by the model. Conserved regions may indicate ones which the model deemed crucial for the overall structure of the Ab. Conversely, if such consensus is not exhibited, we can posit that the model uses different criteria for each species. This could also illustrate which parts of the species’ germline repertoires the model are suitable frameworks for novel CDR sequences.

To investigate this, we generated sequences for the same set of CDRs using different species tags, to see which parts of the sequence change as response to the species.

### Humanization of camelid Abs

Nanobodies can be preferrable over traditional antibodies for their smaller size, higher stability, better tissue penetration, and easier production. Therefore, our ability to humanise them is crucial to the utility of our tool. Since nanobodies differ from Abs in the crucial CDR3 region in length and structure, this task may be far more difficult for our model.

We generate sequences using alpaca CDRs tagged with human identifiers, and validate them based on their T20 score.

## Results

The following section details the results of the experiments we have performed to validate our model.

### Multi-species generation properties

In Figure 2 we plot the KDE of a 2-dimensional t-SNE projection of the original and *de novo* test sequences. When we compare the two right-hand graphs we can clearly see that the biggest differences in the sequences are found in canine species, indicating that canine sequences are the most problematic for our model to generate.

In Figure 1 we plot the difference in the CAI for each of the species between the *de novo* sequences generated based on the test set and the test set sequences. We see that the model keeps the original distribution very well, particularly for murine sequences, with the canine sequences performing the worst. The UAA, UAG and UGA stop codons have been excluded since these do not occur in the natural data at all. This means their CAI for each codon is trivially 0. The small number of malformed sequences in the *de novo* data with stop codons results in large spikes in CAI for those codons.

**Figure 1.**
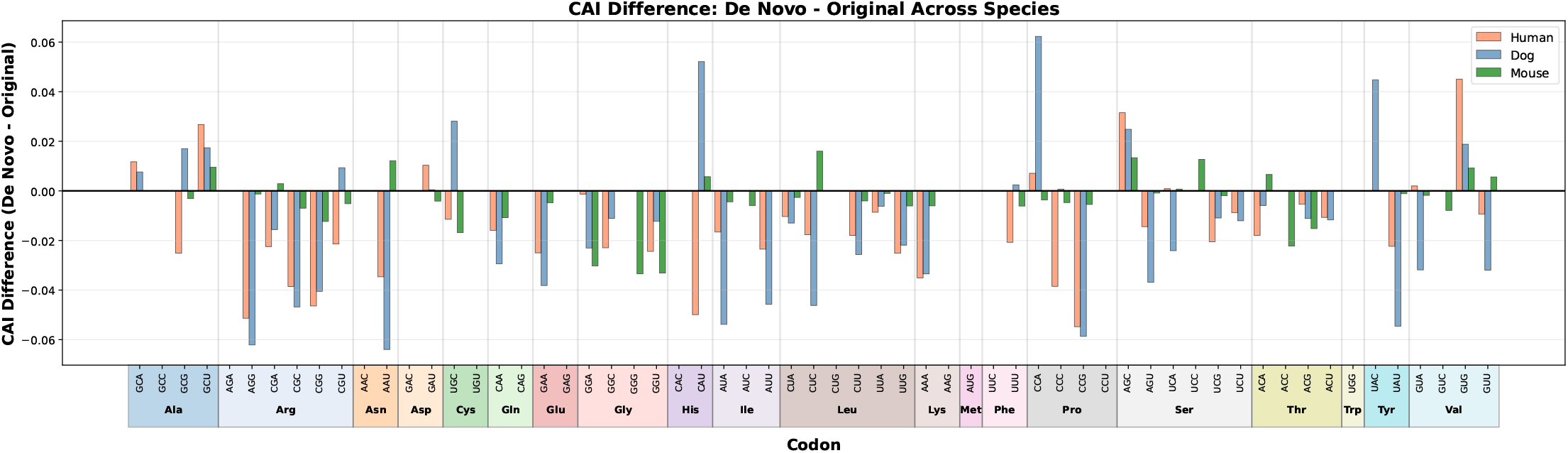
A plot showing the difference in codon adaptation between the *de novo* Ab sequences based on the test set CDRs, and their corresponding natural sequences in the test set. Note that the UAA, UAG and UGA codons are stop codons which are absent from the training data and thus have been left out of the plot.

**Figure 2.**
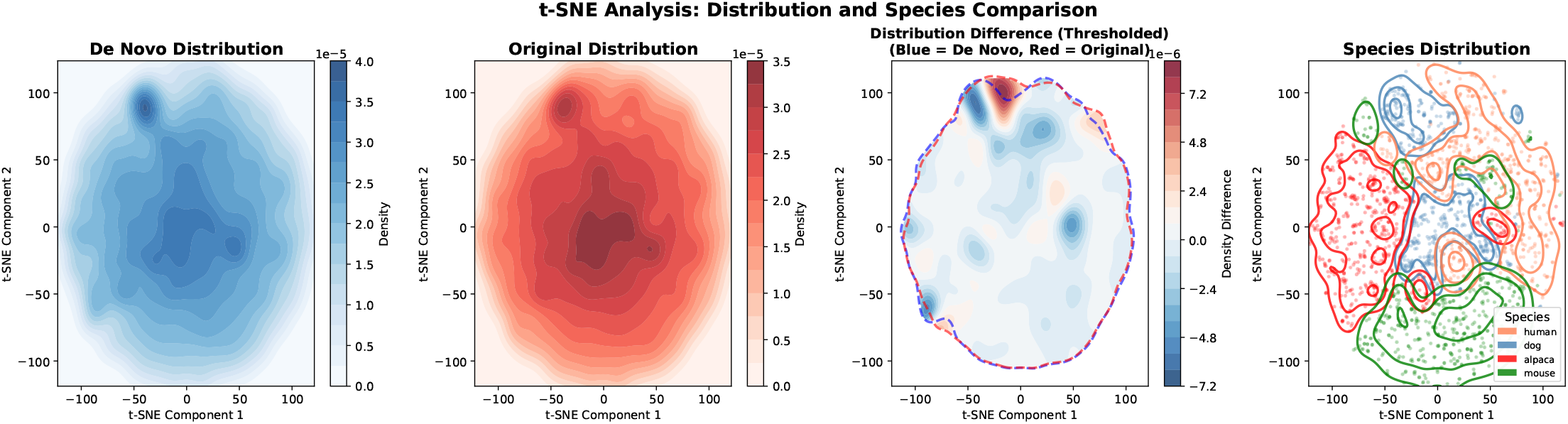
KDE plots of the t-SNE values of the distributions of the one-hot encoded test set sequences. All plots have been calculated using the same t-SNE process. The De Novo Distribution plot shows the KDE heatmap of the *de novo* generated sequences. The Original Distribution plot shows the KDE heatmap of the original sequences used to generate the *de novo* sequences. The Distribution Difference plot shows the difference in the two distributions in their high density regions. The high density regions, represented by the blue and red outlines for the *de novo* and original sequences respectively, are those with a KDE density in the top 40% of the distribution. The colours represent the difference between the distributions, such that blue indicates sequences over-represented in the *de novo* set, and red in the orignal set. The Species Distribution plot shows the KDE outlines and points of the sequences, grouped by species.

Overall, excluding the stop codons, we achieve a mean absolute CAI difference of 0.0148 for humans, 0.0191 for dogs and 0.006 for mice.

We then compared the reconstruction error for the multi-species and mono-species models using the *E*_*B*_ between the *de novo* sequences and the original sequences. We also measure their humanness and canineness using the T20 score [7] for human sequences and cT20 score for the canine sequences [8]. We furthermore score humanness using the OASis (Observed Antibody Space identity score) metric using the relaxed setting [16]. This can be seen in Table 2. In both cases, the differences in performance are relatively minor. As expected, the mono-species model performs slightly better for the canine sequences. This is expected since even with the rebalancing, the multi-species model might put too much emphasis on the human data. Interestingly, the multi-species model seems to perform better on human data, showing that the diverse data helped improve its performance on the majority species.

**Table 2.** Comparison of the results of the multi-species model (i.e. finetuned only on data from all 6 species) and the mono-species models (i.e. finetuned only on human or canine data respectively). Hu stands for human, Ca for canine. *E*_*B*_(*N*) and *E*_*B*_(*P*) stand for the nucleotide and protein level biological loss, as defined in Equation 2. (c) T20 stands for the T20 humanness and cT20 canineness measures as appropriate. OASis is the Observed Antibody Space identity search identity humanness metric, using the relaxed setting [16].

|  | Multi-species Model |  |  |  | Mono-species Model |  |  |  |
| --- | --- | --- | --- | --- | --- | --- | --- | --- |
| | $E_B(N)$ | $E_B(P)$ | (c)T20 | OASis | $E_B(N)$ | $E_B(P)$ | (c)T20 | OASis |
| Hu | 13.05 | 69.99 | 0.9492 | 0.9191 | 13.39 | 71.40 | 0.9509 | 0.9178 |
| Ca | 20.10 | 88.79 | 0.9850 |  | 27.53 | 118.90 | 0.9891 |  |

When we compare the outputs of the multi-species model to those of the mono-species model, we find that about 69% of the *de novo* sequences are are exactly the same between the mono- and multi-species models, but conversely 0% are the same for the canine models. When taken together with the results from subsection 3.3, this can potentially be explained by the mono-species model reusing more of the germline canine-repertoire than the multi-species, which learns a more complete, interlingual representation of FRs which are then adjusted towards the species. This effect would be less pronounced in the human data, since it spans a much larger portion of the dataset.

### High-throughput generation

Generating using a top-*k* sampler with *k* = 10, we generate 10,000 sequences single CDR sequence, without changing the target species. After generating 10,000 sequences, we computed the number of mutations between each *de novo* sequence and the natural sequence as well as the number of unique sequences. A summary of those can be found in Figure 3. We see that there is significant diversity even among the sequences highly similar to the original, with 3,354 unique candidates with at most nine mutations. Ab properties like expression and binding affinity are still impossible to accurately predict *in silico*, so a diverse set of candidate sequences with high similarity to the target is beneficial for further clinical use.

**Figure 3.**
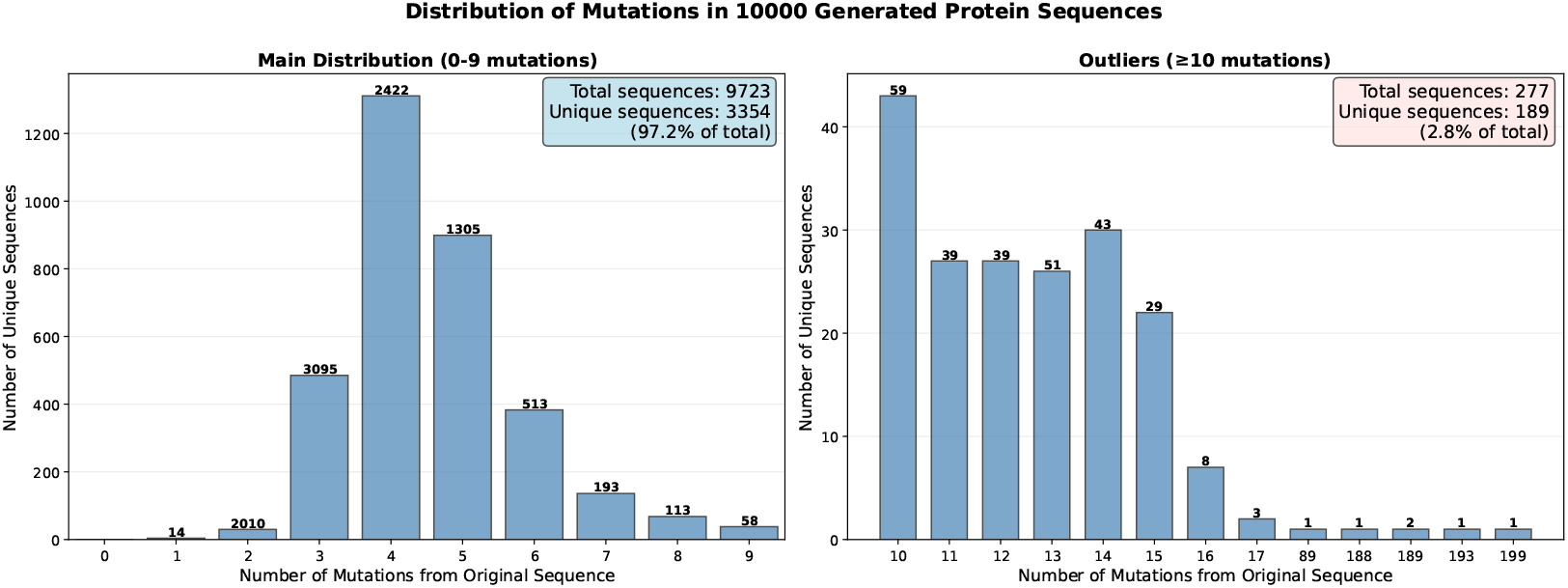
A histogram showing a summary of the 10,000 sequences generated for a single CDR input. These sequences were generated using top-*k* sampling (*k* = 10). The bars correspond to the number of unique sequences generated with a particular number of mutations compared to the original sequence with the numbers above each bar corresponding to the total number of sequences in each bar. The outliers with more than 9 mutations are plotted separately in the right-hand graph to better show the distribution of the low mutation sequences in the left-hand graph.

### Interlingual Ab representation

To try to gain understanding of the underlying representation of Abs present in our model, we compare the model outputs for the same CDRs but with different species tags. Two sets of CDRs are chosen randomly from the test set, one murine and one human. We generate one sequence for each set of CDRs for each of the species used in the project as well as one without a species tag. These sequence-species harmonised sequences are aligned with their respective original sequence using multiple sequence alignment with the MEGA suite’s muscle alignment algorithm [6]. The sequence consensus can be found in Figure 4. We can see that, as expected, in both cases the *de novo* sequence matching the species has the fewest mutations compared to the original.

**Figure 4.**
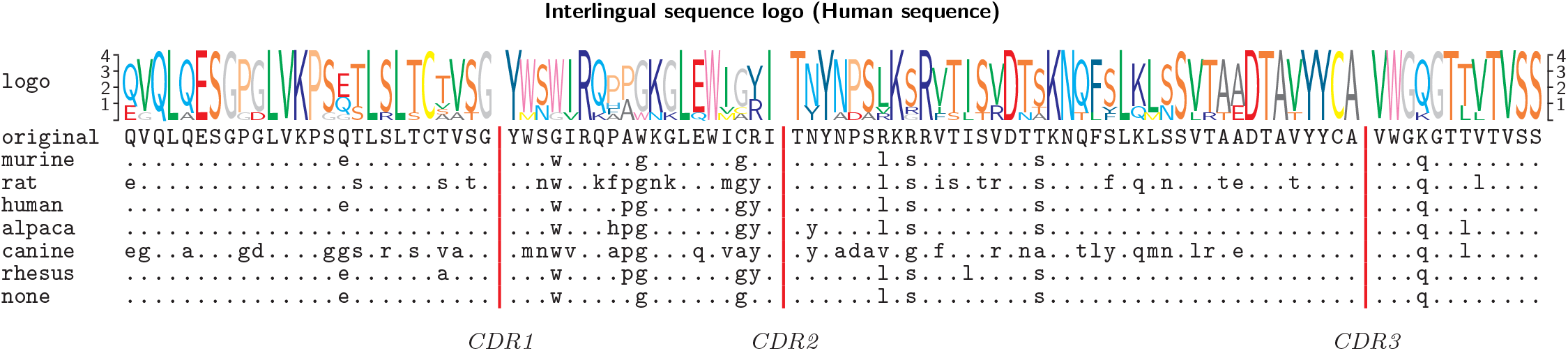
Consensus sequence alignment for a germline human sequences to seven *de novo* sequences based on its germline CDRs. Each of the seven is generated with a different species token. The dots represent a match with the original sequence, letters represent mutations. CDRs are hidden, since they are not affected during generation. On the top is a sequence logo representing the conservation of residues at each position.

All the *de novo* generated sequences share the majority of their sequences among one another, and differ in only a few mutations to adapt them towards their respective species. In fact, in most cases, the sequence-species harmonised sequences differ only by a few mutations from the species-free sequence. The untagged sequence seems to contain all the most common amino acids from the other species. We can see this sequence as an interlingual representation of what the structure of the CDRs demands, which then is mutated to fit the individual species’ needs.

In the case of the murine-based sequence the mouse, rat, canine and monkey *de novo* sequences, and in the case of the human-based sequence the mouse and monkey *de novo* sequences are identical to the species-free sequence. This might indicate that these are already sufficiently close to the interlingual representation, or conversely, that the model has not learned sufficiently how to make sequences for those species. The latter is unlikely due to the large proportion of murine data.

### Humanization of camelid Abs

After generating *de novo* sequences from alpaca CDRs tagged with human identifiers, we compare their human-likeness to the original sequences using the T20 metric. The average T20 score of the sequences when comparing FRs only goes from an average of 0.777 for the original alpaca sequences, to an average of 0.810 for the humanised sequences. This increase brings it barely above the 0.80 score cut-off proposed in the original T20 paper. While this is enough, this is modest compared to the 0.9492 achieved for *de novo* sequences generated from human CDRs. This modest performance can be explained by the structural differences between human and alpaca Abs, such as the significant difference between their CDR-3 lengths. While the T20 score is suboptimal, it may represent an optimal balance between the humanness of the sequence and preserving the tertiary structure of the CDRs.

## Discussion

We have shown our model is capable of producing sequences of a highly similar distribution to the training data. Our model’s outputs match the distributions of natural sequences of each species in terms of codon adaptation, sequence distribution as well as T20 and cT20 metrics for humans and dogs respectively. Notably, our model achieves over 90% OASis identity score, outperforming the Sapiens tool, which scores below 80% [16].

We have demonstrated that the model is able to generate sequences from almost all parts of the natural distribution, rather than repetitively selecting germline FRs which have many associated CDRs. A notable exception are canine sequences. The protein level performance of our multi-species model is slightly worse than that of DoggifAI, which reached an *E*_*B*_(*P*) = 74.90 using the same data [8]. Notably, this is also much better than the analogously trained mono-species model. This shows the added challenge of working in the mRNA space. Another explanation may be that the error metrics *E*_*B*_ and *E*_*d*_ do not take into account the many-to-many relationship between sets of CDRs and sets of FRs; the same set of CDRs can be found with several different sets of FRs and vice-versa. This means that these metrics by definition cannot reach 0, since there are several known biologically correct answers it can choose from. Future projects should take into account this many-to-many relationship between CDRs and FRs, for example by calculating metrics towards all of the valid sets of FRs and selecting the lowest. Though this fell outside of the scope of this computational work, experiments should be done measuring desired properties like binding affinity through *in vitro* validation as was done in HuA bDiffusion [11].

Nevertheless, the ability to jointly optimise the protein sequence and codon distribution is a key benefit of working with mRNA data. It allows the model to balance retaining the secondary structure and resultant binding affinity, the CAI needed for optimal vector expression, and species harmonisation of the protein sequence.

The multi-linguality of our more also allowed us to investigate its understanding of Ab structure in more detail. We see that the model does not treat every species as an entirely different class of problem, and its outputs are similar for each species. Instead of generating entirely dissimilar sequences, we can observe that our model adapts Abs by adding species-specific mutations to an underlying interlingual Ab sequence.

The transformer’s left-to-right generation of sequences may not be optimal for modelling the complex, long-range interactions of amino acid chains. Decisions made at the start of the generation enforce decisions later in the generation, and there is no way for the model to make changes in already generated tokens.

One potential approach to address this limitation is to use diffusion models [21] instead. These models generate their outputs iteratively which might make them more suitable for generating coherent protein sequences. Such models are already being successfully applied to protein level humanisation, showing their promise for mRNA level multi-species sequence-harmonisation [11].

The performance of our model in terms of humanisation of alpaca Abs is modest in terms of the T20 metric. T20 might be a flawed metric to use in this particular instance, since it relies on similarity to an existing database of human heavy chain Ab FR sequences. Human FRs appropriate for the longer CDR3 regions might not exist. Thus even if the sequences were humanised properly, the T20 score might still be low. Furthermore, it remains an open question what degree of humanisation, if any, is needed for safe Nb therapeutics. In fact it has been observed that nanobodies present only a minor immunogenicity risk in humans [2]. Nevertheless, even minor risks need to be addressed for a widespread adoption of this therapeutic class. Here too, *in vitro* validation is the only certain way of assessing the putative decrease of immunogenicity achieved by our tool.

## Author contributions statement

**D.G**.: Writing – original draft, visualisation, validation, software, methodology, investigation, formal analysis, data curation, conceptualisation; **M.K**.: Writing – original draft, writing – review & editing, resources, investigation, data curation; **M.P**.: conceptualisation; **S.C**.: Writing – review & editing, supervision, resources, project administration, conceptualisation; **J.A.A**.: Writing – review & editing, supervision, resources, project administration, conceptualisation.

## Acknowledgments

This work was supported by the United Kingdom Research and Innovation (grant EP/S02431X/1), UKRI Centre for Doctoral Training in Biomedical AI at the University of Edinburgh, School of Informatics and the Royal Academy of Engineering UK Intelligence Community Postdoctoral Research fellowship grant number (ICRF2122-5-133). We appreciate the provision of compute resources through Isambard AI (University of Bristol/UKRI).

